# Spatially resolved DNP-assisted NMR illuminates the conformational ensemble of α-synuclein in intact viable cells

**DOI:** 10.1101/2023.10.24.563877

**Authors:** Jaka Kragelj, Rupam Ghosh, Yiling Xiao, Rania Dumarieh, Dominique Lagasca, Sakshi Krishna, Kendra K. Frederick

**Affiliations:** Department of Biophysics, UT Southwestern Medical Center, Dallas, TX 75390-8816; Center for Alzheimer’s and Neurodegenerative Disease, UT Southwestern Medical Center, Dallas, TX 75390

**Keywords:** DNP solid-state NMR, in-cell NMR, a-synuclein, DNP methods

## Abstract

The protein α-syn adopts a wide variety of conformations including an intrinsically disordered monomeric form and an α-helical rich membrane-associated form that is thought to play an important role in cellular membrane processes. However, despite the high affinity of α-syn for membranes, evidence that the α-helical form is adopted inside cells has been indirect. DNP-assisted solid state NMR on frozen cellular samples can report on protein conformations inside cells. Moreover, by controlling the distribution of the DNP polarization agent throughout the cellular biomass, such experiments can provide quantitative information upon the entire structural ensemble or provide information about spatially resolved sub-populations. Using DNP-assisted magic angle spinning (MAS) NMR we establish that purified α-syn in the membrane-associated and intrinsically disordered forms have distinguishable spectra. We then introduced isotopically labeled monomeric α-syn into cells. When the DNP polarization agent is dispersed homogenously throughout the cell, we found that a minority of the α-syn inside cells adopted a highly α-helical rich conformation. When the DNP polarization agent is peripherally localized, we found that the α-helical rich conformation predominates. Thus, we provide direct evidence that α-helix rich conformations of α-syn are adopted near the cellular periphery inside cells under physiological conditions. Moreover, we demonstrate how selectively altering the spatial distribution of the DNP polarization agent can be a powerful tool to observe spatially distinct structural ensembles. This approach paves the way for more nuanced investigations into the conformations that proteins adopt in different areas of the cell.

## INTRODUCTION

In-cell structural biology enables the study of protein conformation in environments that maintain the identity, stoichiometry, concentrations and organization of the myriad of biomolecules that can interact with a protein of interest.(1-4) Nuclear magnetic resonance (NMR) is uniquely suited to study proteins in these complicated contexts.(2, 5-8) NMR has the resolution and specificity to study atomic-level protein conformations of isotopically-labeled proteins in complex environments that contain molecules with a wide range of molecular sizes and dynamic properties.

Both solution and solid states NMR are well-suited to investigate molecules inside cells and can provide highly complementary insights. Solution state NMR excels in the characterization of small, dynamic biomolecules. However, it is less well-suited for larger, slower moving biomolecules because the formation of protein complexes and/or interactions with membranes results in signal attenuation. In contrast, magic angle spinning (MAS) solid state NMR, particularly when performed under cryogenic conditions, can report directly on the entire conformational ensemble – including protein-protein complexes and proteins that associate with membranes(9-11). While solid state NMR has historically been limited by experimental sensitivity, with the sensitivity gains conferred by dynamic nuclear polarization (DNP), solid state NMR has the sensitivity to detect proteins at their endogenous concentrations(1, 12-16). Thus, DNP-assisted MAS solid state NMR is particularly well-suited to directly investigate the conformations of proteins in cellular settings because it can report directly on the ensemble of sampled protein conformations, including the conformations for proteins in complex with membranes or other cellular constituents. We recently developed protocols for in cell DNP-assisted NMR that result in efficient DNP enhancements and are compatible with cellular viability(17-20).

Because DNP increases the sensitivity of NMR spectroscopy through the transfer of the large spin polarization of an unpaired electron to nearby nuclei(21), the sensitivity enhancements from DNP rely upon proximity to the polarization agent.(22) Thus, DNP-enhanced MAS NMR experiments are biased towards observation of molecules that are accessible to polarization agents. These polarization agents are typically introduced into a sample by doping with millimolar concentrations of stable biological radicals (23-25). In our recent work describing methods for DNP MAS NMR on intact viable cells (19, 26), we examined two of many potential approaches to deliver polarization agents to intact cells. In that work, we introduced the polarization agent, AMUPol (25), to cells by electroporation of intact cells in the presence of AMUPol and by incubation of intact cells with AMUPol and compared the distribution of the polarization agent throughout the cellular biomass for cells (26). AMUPol was homogenously distributed inside cells when it had been introduced by electroporation. Thus, data from experiments on such samples report quantitatively on the entire structural ensemble. In contrast, AMUPol was inhomogeneously distributed in cells when it was delivered by incubation; the signal intensity from DNA in the nucleus was lower than the signal intensity from proteins and RNA in the cytoplasm. Thus, data from experiments on such samples report qualitatively, not quantitatively, on the structural ensemble. Any observed conformation in such samples exists, but the population of that conformation relative to any other cannot be determined directly from integrated peak intensities. Collectively, this work indicated DNP-assisted MAS NMR could be used both to understand the conformational ensemble of a protein and to uncover spatial biases in the distribution conformations of a protein throughout a cell.

The protein α-syn adopts a wide variety of conformations which include an α-helical rich membrane-associated form that is thought to play an important role in cellular membrane processes(27) and an intrinsically disordered monomeric form that is the dominant form in solution (28-30). Purified α-syn can be introduced into cultured mammalian cells by electroporation and remains uniformly distributed throughout the cell for days (31). Initial solution state NMR studies of α-syn inside cells indicated that it was a compact intrinsically disordered monomer (31). Surprisingly, despite the high affinity of α-syn for membranes in purified settings (32), there was no evidence of membrane-associated α-syn inside cells (31). A subsequent in-cell solution state NMR investigation of α-syn found that the signal attenuation observed at the first 12 amino acids of α-syn resulted from transient interactions of α-syn inside cells with chaperone proteins (2). Moreover, they found that chaperone association and membrane-binding of α-syn are mutually exclusive in purified settings. Reduction of the cellular chaperone levels resulted in co-localization of α-syn with mitochondria and signal attenuation of the first 90 amino acids of α-syn inside cells(2), a signature of membrane-associated α-syn (33-35). While this strongly suggests that α-syn can adopt an α-helical rich membrane-associated form inside cells, α-helical rich conformations were not directly observed; their presence was only inferred from the pattern of signal attenuation. Moreover, the signal attenuation pattern was only observed in the setting of a cell with a highly perturbed chaperone environment. Thus, the conformations that α-syn adopts in cells remain poorly understood, and the α-helical rich form, which is thought to be the functional conformation, has not yet been directly observed in unperturbed cells.

Despite extensive study of α-syn, fundamental questions remain about its conformational distribution in healthy cells, particularly regarding its membrane-associated α-helical form. In this work, we introduce α-syn to HEK293 cells, freeze the cells and directly assess the entire conformational ensemble using DNP-assisted MAS NMR. Using purified samples, we find that the membrane-associated α-helical rich conformation and the intrinsically disordered conformation are easily distinguished by their chemical shifts under DNP conditions. Using these two spectra, we model the conformational ensemble of α-syn inside cells as a linear combination of the membrane-associated α-helical rich conformation and intrinsically disordered conformation and quantify the population of α-helical rich α-syn present in healthy cells.

Furthermore, by controlling the spatial distribution of the polarization agent, we determine whether the distribution of α-syn conformations is uniform throughout the cell or if the distribution of conformations has a spatial bias.

## RESULTS

To determine if the α-helical rich nanodisc-associated form and the frozen intrinsically disordered monomeric form of α-syn could be distinguished under the experimental conditions required for efficient DNP-enhanced experiments, we collected two-dimensional ^13^C-^13^C correlation spectra of uniformly ^13^C labeled α-syn associated with nanodiscs and frozen in solution. To do so, we compared the peak shapes and centers for the glycine C^α^-C’, non-glycine carbonyl C^α^-C’, alanine C^α^-C^β^ and threonine C^β^-C^γ^ regions because cross peaks for those sites are distinguished by 10 ppm or more from other cross peaks. For nanodisc-associated α-syn, the peak centers of these regions were consistent with α-helical chemical shift value (Figure 1, red; Table S1). For frozen monomeric α-syn, as expected(36), these regions had composite peaks that spanned a wide range of chemical shifts (Figure 1, blue), and the central values of these broad peaks were consistent with random coil values (Table S1). The peak centers of the α-helical-rich nanodisc-associated form of α-syn and the frozen intrinsically disordered monomeric forms of α-syn differ by an average of 2 ppm. These two forms are distinguishable by ^13^C-^13^C correlation spectroscopy by both peak center and shape under DNP conditions (Figure 1, left column).

**Figure 1:**
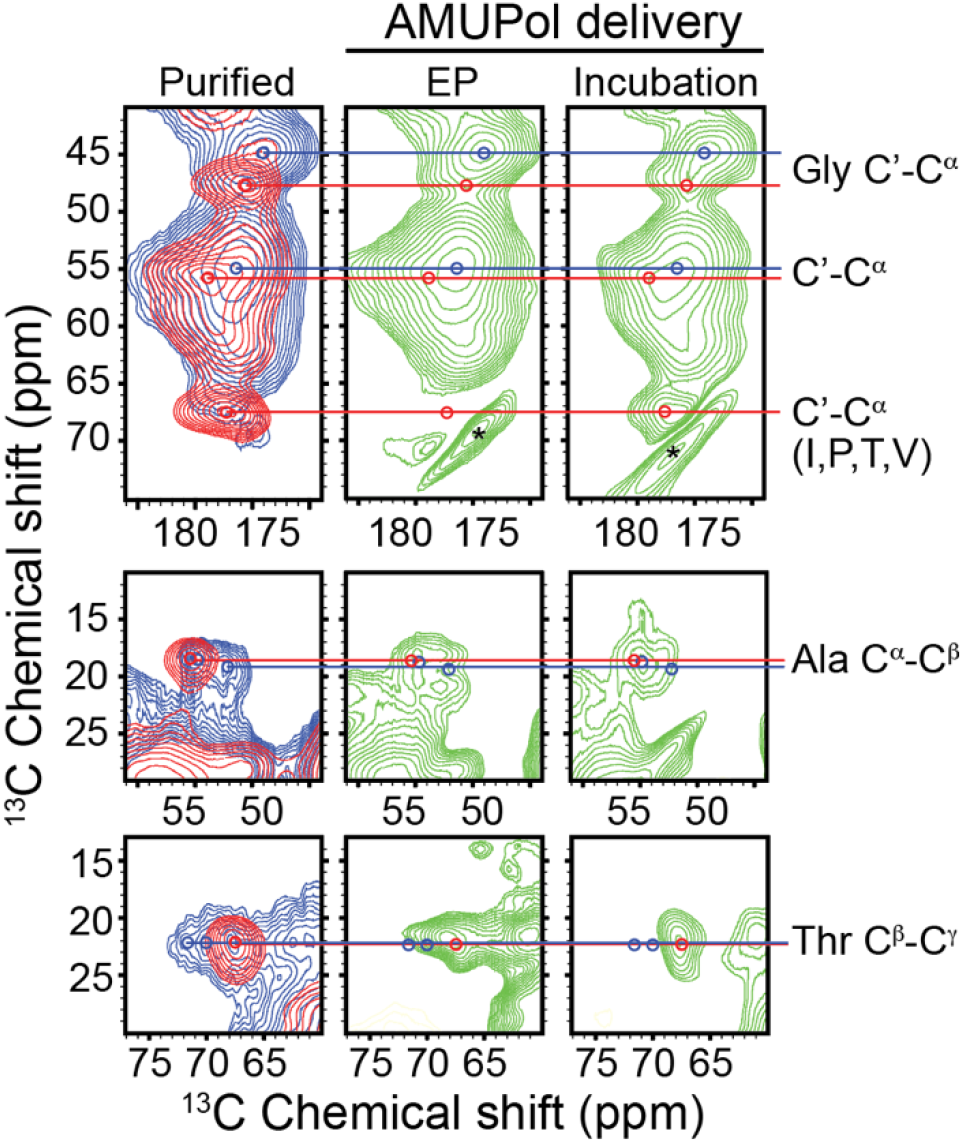
Purified samples of uniformly ^13^C labeled α-syn in the nanodisc-associated form (red) and the frozen intrinsically disordered form (blue) were distinguished by ^13^C-^13^C correlation spectroscopy (DARR, 20 ms mixing) under DNP conditions (left column). The spectra of 75 µM ^13^C labeled α-syn inside intact HEK293 cells where the polarization agent AMUPol was introduced by electroporation (center column) resembles the spectra of the purified intrinsically disordered monomer while the spectra of 75 µM a-syn inside cells where AMUPol was introduced by incubation in 10 mM AMUPol (right column) shares features with the nanodisc associated form. Peak centers for the nanodisc-associated form are annotated with a red circle and peak centers for the frozen monomer are annotated with a blue circle. Spinning side bands are marked with an *. All spectra were recorded at 600 MHz with 12 kHz MAS at 104 K.

To assess the structural ensemble of α-syn inside HEK293 cells, we introduced uniformly ^13^C labeled α-syn at an intracellular concentration of 75 µM using established protocols (Figure S1). This concentration, which is double the endogenous concentration of α-syn in neurons and similar to concentration of chaperone proteins in cells (37), did not result in puncta formation during the experiment (Figure 4). To obtain spatially unbiased spectra, we delivered the polarization agent AMUPol by electroporation, ensuring a homogenous distribution throughout the cellular biomass(26). The resulting ^13^C-^13^C correlation spectra had broad peaks with shapes and centers for the glycine C^α^-C’, non-glycine carbonyl C^α^-C’, alanine C^α^-C^β^ and threonine C^β^-C^γ^ regions that were most consistent with a frozen region of intrinsic disorder (Figure 1, middle column, Table S1). While these data cannot definitively rule out the presence or absence of α-helical rich form of α-syn, they indicate that α-syn predominantly adopts an intrinsically disordered conformation in cells.

To investigate potential spatial bias in α-syn conformations, we next examined cells where AMUPol was delivered by incubation rather than electroporation, a method that produces heterogeneous distribution favoring the cellular periphery over nuclear regions (26). The ^13^C-^13^C correlation spectra for these samples had chemical shifts that were consistent with those of both α-helical and intrinsically disordered conformations (38) (Figure 1, right column, Table S1). For example, the glycine and backbone C’-C^α^ region had maxima with peak centers consistent with those of both nanodisc-associated and intrinsically disordered α-syn (Figure 1, right column). These results indicate that α-syn adopts at least two distinct conformations inside HEK293 cells; an α-helical rich form and an intrinsically disordered form. Moreover, the strong α-helical chemical shifts in this sample suggest that the α-helical rich form is preferentially adopted in the peripheral regions of the cytoplasm.

To better visualize and quantify the structural ensemble of α-syn inside cells, we developed a more specific reporter of α-helical α-syn conformation. Threonine residues are uniformly distributed throughout the region that has α-helical propensity when associated with nanodiscs, are absent from the amino terminal chaperone interaction region and the disordered carboxy terminal region (Figure 2B), and have Cα chemical shift values that are distinct from those of most other amino acids. Thus, we specifically isotopically labeled the threonine residues in α-syn. This approach resulted in α-syn that was ^13^C and ^15^N labeled at the threonine residues with 10% scrambling of ^13^C to glycine but no scrambling to other amino acids. We collected 1D ^15^N-filtered ^13^C spectra of α-syn associated with nanodiscs and frozen in solution. The nanodisc-associated form had a major narrow peak centered near the database value for threonine C^α^ at 68.3 ppm and two minor peaks; one centered at the random coil value for threonine C^α^ and one centered at the α-helical chemical shift of glycine C^α^ (Figure 2, red). The frozen intrinsically disordered monomeric form had a broad major peak that covered the range of threonine C^α^ chemical shift values with a maximum at the random coil value and a minor peak centered at the random coil value for glycine C^α^ (Figure 2, blue). The nanodisc-associated and intrinsically disordered conformations of α-syn were distinguished by a difference in the chemical shift of the major peaks by 7.7 ppm and of the peak width of the major peaks by 4.7 ppm. Thus, these two conformations are easily distinguished by 1D spectroscopy of specifically threonine labeled α-syn.

**Figure 2:**
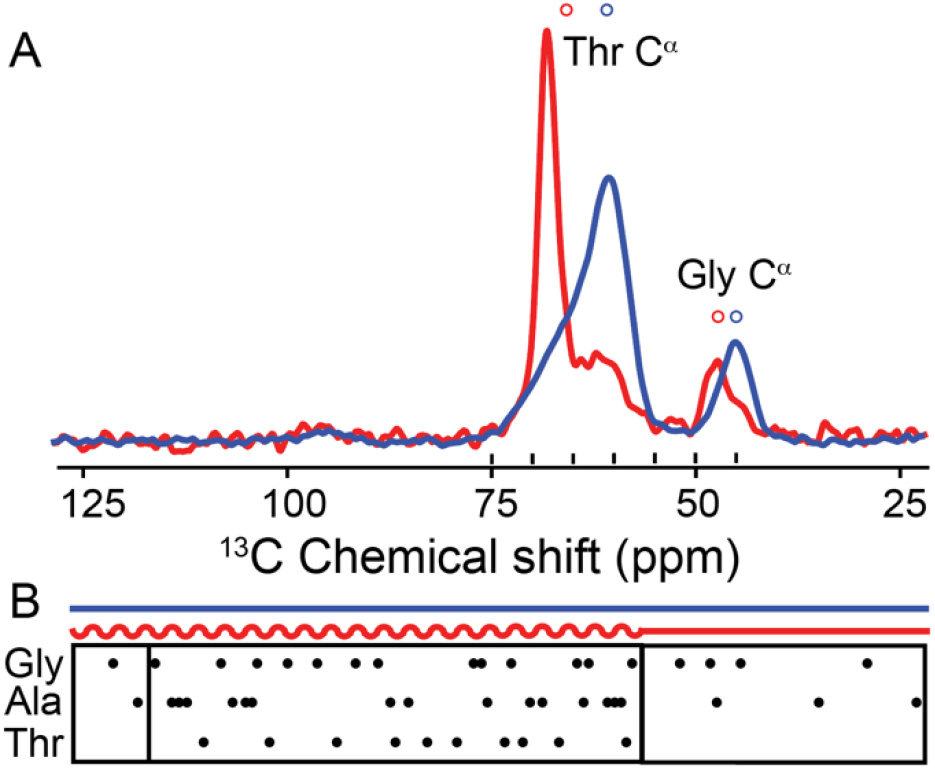
A) α-syn in the nanodisc-associated α-helix rich (red) and frozen monomeric (blue) forms are differentiated by ^15^N-filtered ^13^C spectra collected under DNP conditions on specifically threonine labeled samples. Forward labeling with threonine resulted in ~10% isotope scrambling to glycine. Open circles indicate database chemical shifts for α-helical (red) and random coil (blue) conformations for both threonine and glycine C^α^. B) Cartoon representation of the primary sequence of α-syn with the predicted regions of intrinsic disorder (lines) and α-helices (squiggles) are annotated with the location of glycines, alanines and threonines. Boxes indicate the chaperone binding region for the intrinsically disordered monomeric form (left) and the disordered tail of the membrane associated form of α-syn (right).

To quantify the relative populations of α-helical rich and intrinsically disordered α-syn present inside cells, we collected highly signal averaged 1D ^13^C spectra of threonine labeled α-syn inside cells. We specifically selected for sites in α-syn by using a pulse sequence that selectively detects ^13^C sites within one bond of an ^15^N site (39) providing a 200-fold increase in specificity for labeled α-syn sites relative to natural abundance isotopes in the cellular biomass (22, 40). We then compared ^15^N-filtered ^13^C spectra from HEK293 cells containing 75 µM of threonine-labeled α-syn under two conditions: AMUPol delivery by electroporation versus incubation. The delivery method significantly impacted the spectra. Delivery of AMUPol by electroporation resulted in spectra that resembled that of the frozen intrinsically disordered monomer with a small additional feature consistent with nanodisc-associated α-syn. In contrast, delivery of AMUPol by incubation resulted in spectra dominated by the nanodisc-associated α-syn with a minor feature at the random coil value.

To quantify these differences, we fit each spectrum to a linear combination of two reference spectra: purified nanodisc-associated α-syn (highly α-helical) and purified frozen intrinsically disordered α-syn (Figure 2). We found that in cells where AMUPol was introduced by electroporation, α-syn was predominately intrinsically disordered (92% ± 1) but had a small (8% ± 2) α-helical population (*R*^*2*^ = 0.75) (Figure 3A). Conversely, in cells where AMUPol was introduced by incubation, α-syn was mostly α-helical (60 ± 1%) but had a sizable intrinsically disordered population (40 ± 1%) (*R*^*2*^ = 0.80) (Figure 3B). The AMUPol delivery method altered the α-helical population of the spectra by an order of magnitude.

**Figure 3:**
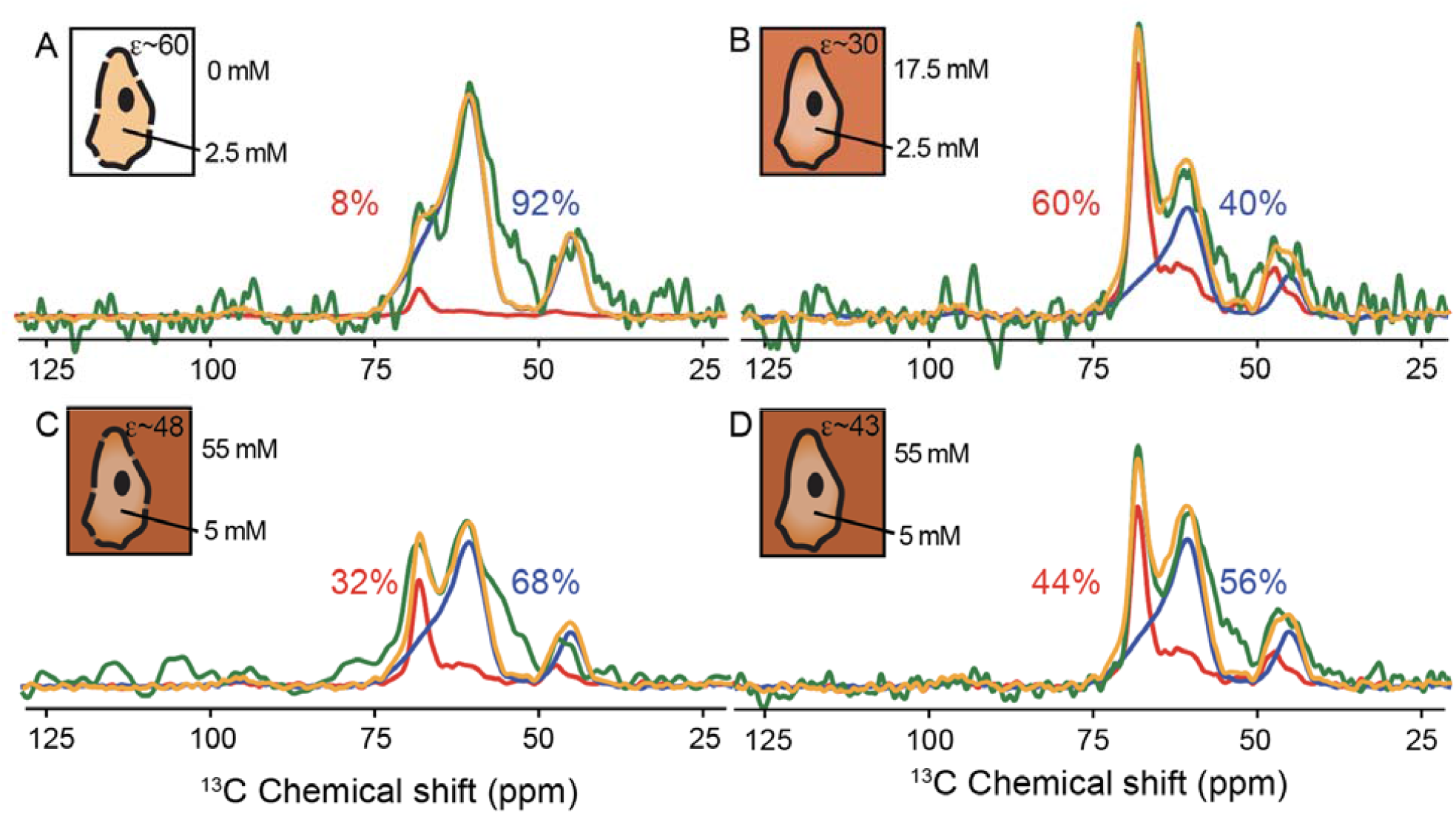
Altering the spatial distribution of AMUPol in cellular samples highlights conformations present in different regions of the cells. A) Delivery of AMUPol via electroporation of cells in the presence of 20 mM AMUPol followed by removal of the extracellular AMUPol prior to data collection reports quantitively on the entire structural ensemble. In contrast, delivery of AMUPol by other methods results in a spatial bias in the resulting spectra. B) Delivery of AMUPol via incubation of cells in 10 mM AMUPol. C) Delivery of AMUPol to cells that had been electroporated in buffer and allowed to recover for 15 minutes before delivery via incubation in 30 mM AMUPol. D) Delivery of AMUPol to cells via incubation in 30 mM AMUPol. The ^15^N-filtered ^13^C spectra of threonine labeled α-syn inside cells (green) is plotted with the spectra of the nanodisc-associated α-helical rich α-syn (red) and the frozen intrinsically disordered monomeric forms (blue) that are scaled by the weighting that resulted in the best fit linear combination (orange). Insets in each panel are cartoon representation of the AMUPol distribution (brown) in the cell and the interstitial space for each AMUPol delivery method. Darker shades represent higher AMUPol concentrations. DNP enhancements (top right corner) and estimated extracellular (top right) and intracellular (bottom right) AMUPol concentrations are annotated. Spectra were recorded at 600 MHz with 12 kHz MAS at 104 K.

**Figure 4:**
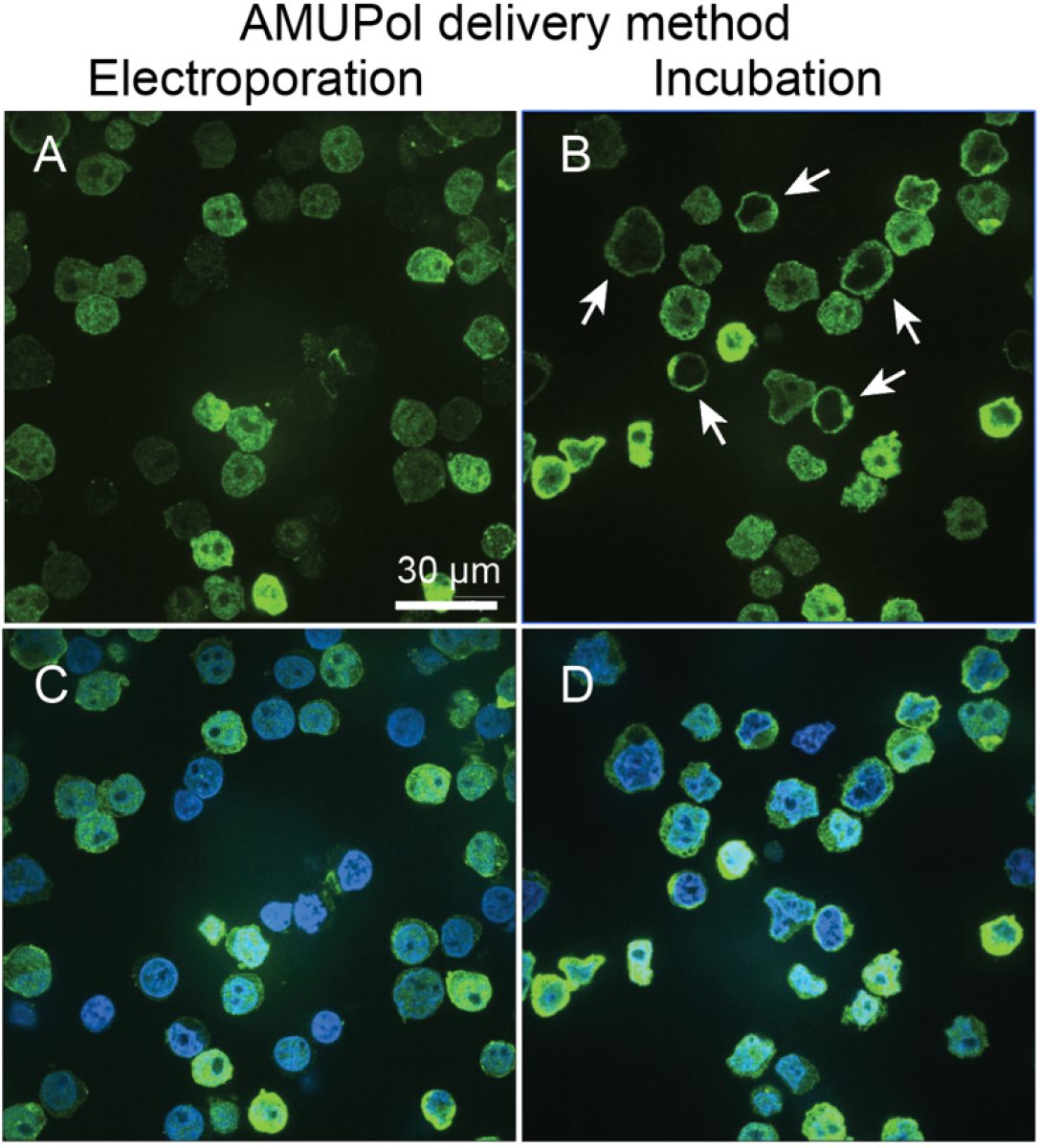
Representative fluorescent microscopy images depicting the distribution of α-syn inside HEK293 cells when AMUPol was delivered by electroporation (A&C) and incubation (B&D). Cells were prepared identically to those used for NMR spectroscopy except cells were fixed and immunostained for α-syn (green) and stained with DAPI (blue). The a-syn was homogenously dispersed throughout most of the cells regardless of AMUPol delivery method, but the a-syn was more concentrated near the cellular periphery (arrowheads) for cells when AMUPol was delivered by incubation.

The order-of-magnitude difference in observed α-helical rich α-syn population between AMUPol delivery methods could arise from three possible sources: changes in the conformational ensemble due to the delivery method, differences in AMUPol distribution throughout the cellular biomass, or a combination of both effects.

To evaluate whether the AMUPol delivery method altered α-syn distribution, we compared α-syn localization in HEK293 cells using an orthogonal method: fluorescent microscopy. Cells were prepared using identical protocols to those for NMR spectroscopy, but were fixed and imaged rather than frozen for NMR analysis. The fixed cells were immunostained with an anti-α-syn antibody and nuclei were visualized with DAPI. When AMUPol was introduced by electroporation, α-syn was predominantly homogeneously dispersed throughout the interior (95.5% of cells, *n* = 223), with only a small fraction showing higher concentration within ~1 µm of the cell periphery (4.5%) (Figure 4). When AMUPol was introduced by incubation, while most cells still showed homogeneous α-syn distribution (80%, *n* = 155), a larger subset displayed peripheral concentration (20%, Figure 4B, arrowheads). Thus, the method of AMUPol delivery altered the localization of α-syn in a sub-population of HEK293 cells, potentially altering the structural ensemble of the sample.

To determine whether this shift in α-syn localization could fully explain the observed NMR spectral changes, we performed a quantitative analysis. Even assuming the extreme case where peripherally-localized α-syn was entirely α-helical, the increased proportion of cells showing peripheral localization (15% increase) could not account for the observed increase in α-helical content (52% increase). Therefore, the dramatic change in α-helical population observed by NMR must result either from the altered AMUPol distribution alone or from a combination of altered AMUPol distribution and changes in the conformational ensemble.

To determine whether the spectral differences arise from AMUPol distribution rather than conformational changes, we modified AMUPol distribution while maintaining the α-syn conformational ensemble. We increased the incubation concentration of AMUPol from 10 mM to 30 mM, which increases intracellular AMUPol concentration (as measured by *T*_*B,on*_ and DNP enhancements) without altering its distribution homogeneity (as measured by the stretch factor β) (26, 41, 42). Assuming cells occupy half of the volume of the rotor, incubation of cells in 10 mM AMUPol results in an intracellular concentration of ~ 2.5 mM and an extracellular concentration of ~17 mM while incubation of cells in 30 mM AMUPol results in intracellular concentrations of ~ 5 mM and an extracellular concentration of ~55 mM AMUPol (26) (Figure 3B,D). Thus, because the degree of inhomogeneity is the same for cells incubated with 10 mM and 30 mM AMUPol, but the concentrations of AMUPol in the intracellular and extracellular space differ, the spatial distribution of AMUPol in these samples must differ. Because intermediate AMUPol concentrations provide optimal DNP enhancements (higher concentrations cause paramagnetic signal attenuation), incubation with 10 mM AMUPol likely produces stronger peripheral enhancement than 30 mM.

The spectral analysis reveals this spatial bias. With delivery of the polarization agent by incubation in 10 mM AMUPol, the spectrum was best fit to 60 ± 1% nanodisc-associated and 40 ± 1% intrinsically disordered conformations (*R*^*2*^ = 0.81) (Figure 3B). In contrast, delivery of the polarization agent by incubation in 30 mM AMUPol resulted in a best fit to 44 ± 1% nanodisc-associated and 56 ± 1% intrinsically disordered conformations (*R*^*2*^ = 0.75) (Figure 3D). This 16% increase in α-helical contribution was observed for the sample with the higher DNP enhancement near the periphery. Thus, α-helical conformations of α-syn predominate near the cell periphery.

To further verify this spatial preference, we modified the AMUPol distribution in electroporated cells. We performed blank electroporation (without AMUPol), allowed cells to recover for 15 minutes, then delivered AMUPol to cells by incubation in 30 mM AMUPol. The resulting AMUPol distribution matched that of a standard delivery by incubation in 30 mM AMUPol (DNP enhancements 47 ± 6, *T*_*B,on*_ 4s ± 1 s, β = 0.82 ± 0.06, *p* > 0.22) and differed significantly from AMUPol delivery by electroporation (*p* < 0.001) (Supplemental Table 2).

The ^15^N-filtered ^13^C spectra of these blank-electroporated cells where AMUPol was delivered incubation was best fit to 32 ± 1% nanodisc-associated and 68 ± 1% intrinsically disordered conformations (*R*^*2*^ = 0.74) (Figure 3B). In contrast, only 8% of the conformational ensemble of α-syn in cells where AMUPol was delivered by electroporation adopts an α-helical conformation. Therefore, the four-fold increase of α-helical conformations observed for electroporated cells when AMUPol was delivered preferentially to the periphery by incubation provides strong evidence for peripheral localization of α-helical conformations. Moreover, because the spatial distribution of AMUPol when it is delivered by incubation isn’t altered when the cells were previously electroporated, we are able to estimate that electroporation reduces the α-helical population of the conformational ensemble by 3%. Therefore, AMUPol delivery by electroporation minimally perturbed the conformational ensemble and the order-of-magnitude difference in observed α-helical rich α-syn population between AMUPol delivery methods is mostly driven by differences in AMUPol distribution throughout the cellular biomass. Thus, we find not only that α-helix rich conformations of α-syn are adopted inside cells under physiological conditions but also that the α-helical rich form is preferentially adopted near the cellular periphery.

## DISCUSSION

Understanding the conformational states of α-syn in cells is crucial for elucidating its biological function. While α-syn is known to adopt both intrinsically disordered and membrane-associated α-helical conformations, direct evidence for its α-helical form inside cells has remained elusive. Here, using DNP-assisted solid-state NMR, we provide the first direct observation of α-helical α-syn in cells. By controlling the distribution of DNP polarization agents throughout the cellular biomass, we were able to both quantify the total conformational ensemble and probe its spatial distribution. Our results reveal that while most α-syn in HEK293 cells exists in an intrinsically disordered state, a distinct minority population adopts an α-helical rich conformation. Moreover, by manipulating the spatial bias of our measurements, we demonstrate that this α-helical form is preferentially localized near the cell periphery, suggesting a potential membrane-associated function.

Previous in cell NMR experiments established that α-syn behaved as a compact intrinsically disordered monomer in various cultured mammalian cell lines, including HEK293 cells (31). This result is consistent with many biophysical and structural experiments of α-syn in both purified and complex biological contexts. Here we found that α-syn in HEK293 cells adopted at least two distinct conformations, one of which is α-helical rich and the other is intrinsically disordered. Because the α-helical form accounts for only 8% of the total ensemble, it is possible that this conformation was present in solution state experiments, but the decrease in peak intensity could not be discerned with confidence above experimental noise (31). However, since our experiments used four times higher α-syn concentrations, the discrepancy may also result from a difference in the underlying conformational ensembles. Nonetheless, we find that α-syn can adopt at least two distinct conformations inside cultured mammalian cells, one of which is rich in α-helices.

While our NMR data clearly demonstrate the presence of an α-helical form of α-syn preferentially localized near the cellular periphery, they do not provide direct evidence for membrane-associated α-syn. Nevertheless, the similarity between our in-cell spectra and nanodisc-associated reference spectra, combined with extensive evidence of α-syn’s membrane interactions in both purified (32) and complex biological settings (43, 44), suggests membrane association. Future experiments using mutant α-syn variants, different isotopic labeling schemes, and additional reference samples could further characterize the specific α-helical rich forms adopted in various cell lines, particularly given α-syn’s selective binding to membranes with specific curvatures (45, 46) and potential formation of α-helical rich tetramers inside cells (47).

The spatial resolution achieved in this study highlights the importance of understanding polarization agent distribution in DNP-assisted MAS NMR experiments (19, 26, 48-50). While fluorescent microscopy of tagged polarization agents can provide spatial information (50), this approach is limited to reduction-resistant agents since fluorescence cannot distinguish between DNP-active and inactive forms. Our use of DNP properties (enhancements, build-up times, and β-factor) to characterize polarization agent distribution offers advantages, as these properties specifically report on biomolecules near DNP-active molecules (26). This approach proves particularly valuable in cellular settings, where the inhomogeneous distribution of AMUPol results from radical reduction kinetics rather than membrane permeability (19, 20, 26).

This study demonstrates how controlling polarization agent distribution can provide spatially resolved structural insights in intact viable cells. Building on recent work examining nuclear proteins using isolated nuclei (50), our approach achieves subcellular specificity by manipulating polarization agent distribution in whole cells. The development of biostable polarization agents (50-52) and targetable polarization agents could further enhance this capability, opening new possibilities for investigating protein conformations within specific cellular compartments. Our finding that α-syn preferentially adopts α-helical rich conformations near the cellular periphery exemplifies the unique insights possible through spatially resolved DNP NMR in intact cells.

## SUPPLEMENTAL INFORMATION

Supplemental information is available for this article.

## ACKNOWLEDGEMENTS

R.G was supported by fellowship from the O’Donnell Brain Institute Neural Science Training Program. D.L. was supported by NIH MB T32 GM008297. S.K. was supported by UT Dallas and the Cecil and Ida Green Foundation via The Green Fellow’s Program. This work was supported by grants from the National Institute of Health [NS111236], the National Science Foundation [1751174], and the Welch Foundation [1-1923-20200401] to K.K.F.

## Methods

### α*-syn expression and purification*

The *E. coli* strain BL21 DE3 was used to express isotopically labeled wild-type α-syn. Uniformly isotopically labeled protein was expressed by growth of cells in 4 L LB to an OD_600_ of 0.6 followed by concentration and resuspension in 1L of M9 media containing 1 g/L of ^15^N chloride and 4 g/L of ^13^C enriched glucose (53). The protocol for specific isotopic labeling of threonines was based on previously published protocols (54, 55). First, the *E. coli* cells were grown in 4 liters of M9 supplemented with natural abundance threonine (50 mg/mL), α-ketobutyrate (100 mg/mL), and natural abundance glycine (500 mg/mL). Cells were grown at 37 °C until the OD_600_ of the culture reached 0.6 – 0.8. The *E. coli* cells were then harvested, washed once with 1x M9 salts (53), and transferred to 4 liters of M9 supplemented with ^13^C,^15^N-threonine (50 mg/ml), α-ketobutyrate (100 mg/ml), and natural abundance glycine (500 mg/ml). Cells were then grown at 37 °C until they reached OD_600_ of 1.5. Expression of α-synuclein was induced with addition of 1 mM IPTG. Cells were harvested after 3 hours of expression at 37 °C.

Purification was performed as described (56). Briefly, cells were frozen after the expression, resuspended in lysis buffer containing a detergent (20 mM Tris pH 8.0, 1 mM EDTA, 0.1% Triton-X) and incubated for 30 minutes at 37 °C. After the incubation, DNA and RNA were digested by adding 2.5 µL Omni Nuclease and 2.5 uL DNAse while supplementing the solution with 10 mM MgCl_2_ and 10 mM CaCl_2_ final concentration. After 1 hour incubation at 37 °C the excess metal was chelated by addition of 5 mM EDTA. Insoluble cell debris was separated from the supernatant by centrifugation at 4,000 x g for 15 minutes. The concentration of NaCl in the supernatant was adjusted by mixing 6 parts of supernatant with 1 part of 5 M NaCl. The supernatant was heated above 90 °C for 10 minutes in a water bath, after which the supernatant was cooled in a room temperature water bath and on ice. The proteins that precipitated during the boiling step were pelleted by centrifugation at 20,000 x g for 20 minutes at 4 °C. The α-synuclein in the soluble supernatant was precipitated by addition of cold saturated ammonium sulfate at 1:1 ratio (final concentration of 50 % saturated ammonium sulfate at 4 °C) and incubation overnight at 4°C with gentle stirring. The precipitated protein was separated from the supernatant by centrifuging at 4000 x g for 20 minutes at 4 °C. The pellet was dissolved in 20 mM Tris pH 8.0, 20 mM NaCl. The solution was loaded onto a Q-sepherose column and α-synuclein was eluted with a salt gradient of 0-500 mM NaCl. The α-synuclein containing fractions were pooled and concentrated to 1 mM and then run over a size exclusion column (Superdex 75 Increase HiScale 26/40, 40 cm). Protein purity was assessed by gel electrophoresis and was determined to be greater than 98%.

### Preparation of frozen intrinsically disordered monomers

The stock α-synuclein solution at 1 mM was diluted in D_2_O to result in 10 mM sodium phosphate buffer at pH 7.0 at a 12:88 ratio of H_2_O:D_2_O. Glycerol was added to a final concentration of 15 % *d*_*8*_-^12^C-glycerol resulting in a 10:75:15 ratio of H_2_O : D_2_O : *d*_*8*_-^12^C-glycerol. AMUPol was dissolved in the sample to result in a final concentration of 10 mM AMUPol. The sample was transferred to a rotor and stored at −80 °C until measurements. Prior to measurement samples were briefly warmed to room temperature to mark and cap the rotor. Room temperature rotors were inserted into the probe that was pre-equilibrated to 100 K. Sample temperature was inferred from the temperature of the stator and decreased from room temperature to 200 K in ~2 minutes followed by a slow decrease to 104 K over 15 minutes.

### Preparation of complex with nanodiscs

The nanodisc scaffold MSP1E3D1 was expressed and purified as previously described (57). POPG (1-palmitoyl-2-oleoyl-sn-glycero-3-phosphoglycerol) was used to assemble the nanodiscs. Buffer used for the nanodisc samples was 20 mM sodium phosphate, 50 mM NaCl, pH 7.4.

Nanodiscs and α-synuclein were mixed at equimolar ratio, and D_2_O was added to obtain an 88:12 D_2_O:H_2_O ratio. The sample was concentrated to the final concentration of 100 μM α-synuclein. Depleted deuterated glycerol (*d*_*8*_-^12^C-glycerol) and AMUPol were added as the last step for a final composition of 15:75:10 for *d*_*8*_-^12^C-glycerol:D_2_O:H_2_O (*v*/*v*/*v*) respectively with 6.8 mM AMUPol.

### Preparation of natural abundance and isotopically enriched HEK293 cells

Human embryonic kidney 293 (HEK293) cells were cultured in Dulbecco’s Modified Eagle Medium (DMEM) supplemented with 10% Fetal Bovine Serum and 1% PenStrep (Gibco). For quantification of the AMUPol distribution through cellular biomass, uniformly isotopically labeled HEK293 cells were cultured in ^13^C, ^15^N labelled media (BioExpress 6000 Mammalian U-^13^C, 98%; U-^15^N, 98%, Cambridge Isotope Laboratories, USA) with 10% (v/v) fetal bovine serum (FBS, qualified, Gibco) and 1% (v/v) PenStrep (Gibco) at 37 °C and 5% CO_2._as previously described (26).

### Introduction of α-syn into HEK293 cells by electroporation

Delivery of α-syn to HEK293 cells by electroporation was performed as described (31). Briefly, confluent adherent HEK293 cells were rinsed with PBS, detached from the culture dish with trypsin/EDTA (0.05 %/ 0.02 %) and collected by centrifugation at 233 x g for 5 min at room temperature. The 100 µL cell pellet was resuspended in 10 mL of PBS and 10 µL of this suspension was mixed with 10 µL of trypan blue (0.4% solution) to determine the number of live cells as assessed by trypan blue membrane permeability with a Countess automated cell counter (Life Technologies) using the manufacturer’s instructions. The cells then were pelleted, washed with electroporation buffer then pelleted again before being mixed with α-syn.

Cells were suspended in 100 mM sodium phosphate, 5 mM KCl, 15 mM MgCl_2_, 15 mM HEPES with freshly added 2 mM ATP, 2 mM reduced glutathione and 900-1000 µM purified α-syn at 40 × 10^6^ cells per mL. 100 µL aliquots (4 × 10^6^ cells) were electroporated with an Amaxa Nucleofector I (Lonza) using the HEK293 pulse sequence. Cells were pulsed twice with gentle mixing followed by a 5 minute room temperature incubation between the two pulses. Ten minutes after electroporation, 0.5 mL of pre-warmed (37 °C) growth medium was added to each cuvette and samples were transferred to a cell culture dish. Cells were returned to the incubator and allowed to recover for 5 h. Once cells regained their adherent morphologies, dishes were washed with PBS and cells were harvested by trypsinization.

### Western blotting

Harvested electroporated cells were counted, pelleted and lysed with RIPA buffer. Samples were fractionated by SDS-PAGE, transferred to polyvinylidene fluoride membrane probed with the anti-a-syn antibody (BD Bioscience 610786). Samples were denatured by incubation at 95 °C for 10 minutes in the presence of 2% SDS before separation. Secondary antibodies were coupled to horseradish peroxidase. Blots were visualized by a standard ECL analysis and band intensities quantified using Image Lab (Bio-Rad).

### Delivery of AMUPol to HEK293 cells

AMUPol was delivered to HEK293 cells by either incubation or electroporation as previously described (26). Briefly, for delivery by incubation, a 50 µL cell pellet was mixed with 50 µL perdeuterated 1x PBS (85% D_2_O + 10% H_2_O, pH 7.4) containing AMUPol (Cortecnet, USA) and 18 µL of *d*_*8*_-glycerol. The 118 µL cell suspension had a final composition of 15% (v/v) *d*_*8*_-glycerol, 75% (v/v) D_2_O and 10% (v/v) H_2_O. For delivery by electroporation, a 50 µL cell pellet was mixed with 100 µL electroporation buffer (SF cell line solution, Lonza) containing AMUPol and electroporated (HEK293 pulse sequence, Lonza 4D-nucleofactor) using manufacturer’s instructions. Post electroporation, cells were allowed to recover for 10 minutes in electroporation buffer containing AMUPol inside the tissue culture hood. Next, cells were washed twice with 50-100 µL (depending on cell pellet volume) of 1x PBS to eliminate the electroporation buffer and any extracellular AMUPol from the sample and the 50 µL cell pellet was resuspended in perdeuterated 1x PBS and *d*_*8*_-glycerol for a final composition of 15% (v/v) *d*_*8*_-glycerol, 75% (v/v) D_2_O and 10% (v/v) H_2_O. For blank electroporation samples, cells were subjected to electroporation in the absence of AMUPol in electroporation buffer, allowed to recover for 10 minutes and then incubated in perdeuterated PBS containing 30 mM AMUPol. For blank electroporation samples, cells were subjected to electroporation in the absence of AMUPol in electroporation buffer, allowed to recover cells for 10 minutes and then incubated in perdeuterated PBS containing 30 mM AMUPol. The DNP matrix had a final composition of 15% (*v/v*) *d*_*8*_-glycerol, 75% (*v/v*) D_2_O and 10% (*v/v*) H_2_O. Cell pellets were transferred to 3.2 mm sapphire rotors by centrifugation, sealed with a silicon plug and then frozen at a controlled rate of 1 °C/min (26).

### Fluorescence microscopy

Cells for fluorescence microscopy were treated identically to those prepared for DNP NMR except that after the manipulation for the introduction of the radical. Cell pellets were fixed in PBS containing 4 % (*w/v*) paraformaldehyde (PFA) for 15 min at room temperature rather than being transferred to rotors and frozen. The cells were washed in PBS twice then permeabilized by incubation in 0.1 % (*v/v*) Triton-X in PBS for 3 min. Cells were pelleted by centrifugation, the supernatant was removed, and the pellet was incubated in fresh PBS for 10 minutes three times before cells were blocked by incubation in 0.1% (*w/v*) BSA (Sigma) in PBS for 1 h. Cells were then incubated for 2 h with primary antibody anti-α-Syn sc-69977 (Santa Cruz, 1:100 dilution) in blocking buffer. Cells were pelleted by centrifugation, the supernatant was removed, and pellet was incubated in fresh PBS for 10 minutes three times before cells were incubated with anti-mouse IgG Alexa-488, (Sigma, 1:1,000 dilution) for 1 h in blocking buffer. Cells were again pelleted by centrifugation, the supernatant was removed, and pellet was incubated in fresh PBS for 10 minutes three times. Suspended cells were allowed to settle onto 44 mm diameter glass bottom dishes (Nunc) for 30 minutes then mounted using ibidi mounting medium containing DAPI. The plates were stored at 4 °C. The cells were imaged by confocal microscopy (Nikon CSU-WI spinning disc confocal) with a 100x objective lens. Excitation and emission were 488/496 nm and 406/460 nm for Alexa488 and DAPI, respectively. Images were analyzed using Fiji software.

### NMR spectroscopy

Rotors were transferred in liquid nitrogen directly into the NMR probe that had been previously equilibrated to 104 K as described (18, 26). All dynamic nuclear polarization magic angle spinning nuclear magnetic resonance (DNP MAS NMR) experiments were performed on a 600 MHz Bruker Ascend DNP NMR spectrometer/7.2 T Cryogen-free gyrotron magnet (Bruker), equipped with a ^1^H, ^13^C, ^15^N triple-resonance, 3.2 mm low temperature (LT) DNP MAS NMR Bruker probe (600 MHz). For ^13^C cross-polarization (CP) MAS experiments, the ^13^C radio frequency (RF) amplitude was fixed at 60 kHz and an ^1^H RF amplitude was 72 kHz. The 90° ^1^H pulse was 100 kHz, the 90° ^13^C pulse was 62.5 kHz, and ^1^H TPPM at 85 kHz for decoupling with phase alternation of ± 15° during acquisition of ^13^C signal. ^13^C-^13^C 2D correlations were measured using 5 ms or 20 ms DARR mixing with the ^1^H amplitude at the MAS frequency. A total of 280 complex points in the indirect dimension were recorded with an increment of 25 μs. DARR experiments were apodized with a Lorenz-to-Gauss window function with IEN-to-GB ratio of 2.5 and the IEN between 20 and 80 for both the *t*_1_ and *t*_2_ time domains. For ^13^C-^15^N 1D spectra either a TEDOR or NCa double CP pulse sequence was used. In the case of TEDOR (39), the mixing time was 1.92 ms with a recycle delay of 3.9 s. For 1D NCa double CP (58) experiments, the contact time was 6 ms with a recycle delay of 3.0 s. The DNP enhancements were determined by comparing 1D ^13^C CP spectra collected with and without microwave irradiation. For *T*_B,on_ measurements, recycle delays ranged from 0.1 s to 300 s. To determine the *T*_B,on_, the dependence of the recycle delay using saturation recovery on both ^13^C peak intensity or volume was fit to the stretched-exponential equation 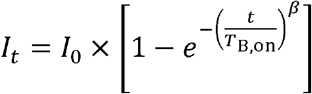.

### Fitting

In-cell spectra were fit to a linear combination of the experimental spectra of nanodisc bound α-syn, which is α-helical, and purified frozen intrinsically disordered α-syn using the generalized least squares regression function in statsmodels.api. The coefficients of the linear regressions were used to weight the monomer and nanodisc-bound data, and a numpy trapezoidal approximation of the integral (trapz) was calculated to determine the relative populations of α-helical and intrinsically disordered α-syn. Code available at https://github.com/kendrakf/Spotlight2023

