## Supplemental Information for "Spatially resolved DNP-assisted NMR illuminates the conformational ensemble of α-synuclein in intact viable cells"

**Current affiliation**:

3. National Institute of Chemistry, Hajdrihova 19, 1001 Ljubljana, Slovenia

*To whom correspondance should be addressed: Kendra K. Frederick

https://orcid.org/0000-0001-6472-8429
https://orcid.org/0000-0002-3095-621X
https://orcid.org/0000-0002-4548-4833
http://orcid.org/0000-0002-4462-7134
https://orcid.org/0000-0003-4404-4366
https://orcid.org/0000-0002-4223-0379
https://orcid.org/0000-0002-1656-5167

| | Glycine | | Alanine | | Threonine | Backbone (no G) | |
|---|---|---|---|---|---|---|---|
| | Cα | C' | Cα | Cβ | Cβ | Cα | C' |
| Frozen monomeric α-syn | 44.5 | 174.2 | 52.3<br>54.4 | 19.2<br>18.3 | 69.5 | 54.7 | 176.4 |
| Nanodisc-associated α-syn | 47.7 | 175.5 | 55.4 | 18.4 | 67.6 | 55.8*<br>67.5^ | 179*<br>177.4^ |
| α-syn inside cells with AMUPol delivered by electroporation | 44.8 | 174.2 | 52.8<br>54.7 | 19.9<br>18.4 | 67.8 | 54.8 | 176.4 |
| α-syn inside cells with AMUPol delivered by incubation | 46.9<br>44.8 | 175.6<br>173.8 | 54.8 | 19.0 | 67.6 | 56.1*<br>67.5^ | 177.8*<br>177.2^ |
| Calculated average random coil | 45.3 | 174.3 | 52.7 | 19.0 | 69.8 | 57.2 | 176.0 |
| Calculated average for α-helix | 47.0 | 176.3 | 54.9 | 18.3 | 68.6 | 58.0*<br>65.8^ | 178.2*<br>177.5^ |

* Peak center for the peak with Cα chemical shifts greater than 50 ppm and less than 65 ppm; excludes glycine as well as isoleucines, prolines, threonines and valines.
^ Peak center for the peak with Cα chemical shifts greater than 65 ppm. Includes only isoleucines, prolines, threonines, and values.

***Table S1**: Peak centers of select sites in α-syn in different environments. For peaks that had two distinguishable maxima, such as the alanine Cα-Cβ peak, the center of the more intense maxima is reported first. Peak centers above and below the diagonal differed by 0.1 ppm or less. Glycine C'-Cα sites are well resolved from other backbone residues, so values are reported separately, and the backbone average excludes glycines.*

*Values for calculated average peak centers are taken from Wang & Jardetzky, 2002. In samples with high a-helical content, the non-glycine C'-Cα peak was resolvable into two major peaks. The peak centers of the two major peaks are reported separately. For a random coil, calculated average chemical shift value was determined by weighting the average chemical shift value for a random coil by the prevalence in the a-syn sequence for all residues except glycines. For α-helices, calculated average chemical shift value was determined by weighting the average chemical shift value for α-helix by the prevalence in the α-syn sequence for both the major peak which excludes glycine, isoleucine, proline, threonine and valine residues as well as the minor peak which includes only isoleucine, proline, threonine and valine residues.*

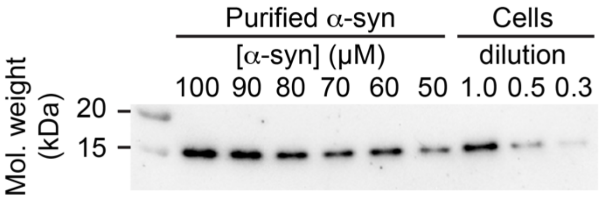


***Figure S1****: Intracellular concentration of α-syn after electroporation was determined by semi-quantitative Western blot. A concentration gradient of purified α-syn served as a standard (lanes 2-7) to establish a calibration curve to determine the concentration of a-syn inside several dilutions of lysed cells (lanes 8-10). The intracellular concentration of α-syn was 75 ± 8 µM.*

| **Protein** | | | | **RNA** | | | | **Lipid** | | | |
|---|---|---|---|---|---|---|---|---|---|---|---|
| ε | $T_{B,on}$ | β | error | ε | $T_{B,on}$ | β | error | ε | $T_{B,on}$ | β | error |
| 42.8 | 3.0 | 0.78 | 1.0% | 52.6 | 3.3 | 0.79 | 1.1% | 31.3 | 3.9 | 0.82 | 1.4% |
| 50.9 | 5.0 | 0.86 | 0.5% | 51.6 | 5.9 | 0.85 | 0.5% | 37.4 | 6.2 | 0.70 | 0.8% |

***Table S2****: Summary of DNP properties for uniformly isotopically enriched HEK293 cells that had electroporated in buffer (a blank electroporation) allowed to recover for 15 minutes and then incubated in 30 mM AMUPol. The DNP enhancement (ε) and $T_{B,on}$ were determined independently for the signals correspond to the protein carbonyl (~176 ppm), nucleotide (~140 ppm), and lipid (~35 ppm) described as defined in Ghosh et al, 2021.*

$T_{B,on}$ is reported in units of seconds and was determined from 13 time points that ranged from 0.5~60 sec. (0.1, 0.5, 1.0, 1.5, 2.5, 3.0, 3.5, 5.0, 7.5, 10, 15, 30, and 60 sec, respectively).

Error is the regression error, determined as reported in the methods section.
